# Functional spatial transcriptomics uncover LMO7 as a fusion-regulated and clinically relevant driver of metastasis in Ewing sarcoma

**DOI:** 10.64898/2026.08.27.747513

**Authors:** Veronika Buršić, Heng Luo, Clémence Henon, Lianghao Mao, Anna C. Ehlers, Angelina Yershova, J-Ann M. Lego, Jing Li, Martha J. Carreño Gonzalez, Richard Arndt, Ana Sastre, Javier Alonso, Uta Dirksen, Wolfgang Hartmann, Ashok Kumar Jayavelu, Moritz Gerstung, Thomas G. P. Grünewald, Florencia Cidre-Aranaz

## Abstract

Metastatic dissemination represents the major determinant of poor clinical outcome across cancer entities. Yet, how driver oncogenes shape transcriptional programs facilitating metastasis is poorly understood. In Ewing sarcoma (EwS) – a highly aggressive pediatric bone and soft-tissue sarcoma driven by chimeric FET::ETS transcription factors – low activity of the fusion oncoproteins is thought to promote metastasis, but the underlying molecular mechanisms remain largely elusive.

Here, using spatially resolved functional transcriptomics in EwS patient tumors, we identify a distinct transcriptional state at the invasive tumor front, that in contrast to the tumor core, is characterized by lower FET::ETS activity and induction of the multifunctional shuttle LIM domain only protein 7 (LMO7). Integrating these data with clinical information reveals that high *LMO7* expression is associated with poor outcomes. Gene network analysis of patient tumors and integrated proteomic and transcriptomic profiling of EwS cell lines following inducible *LMO7* silencing highlight *LMO7* as a central regulatory hub orchestrating epithelial-mesenchymal transition (EMT) and cytoskeletal remodeling in EwS. Functional experiments demonstrate that *LMO7* silencing decreases clonogenicity and migratory capacity *in vitro* and suppresses primary tumor growth and metastatic dissemination *in vivo*.

Collectively, these findings identify LMO7 as a clinically relevant effector of FET::ETS fusions in EwS, and illustrate how integrating functional, spatial and clinical data can uncover oncogene-driven effectors of metastasis.

## INTRODUCTION

Ewing sarcoma (EwS) is a highly aggressive malignant bone and soft-tissue tumor predominantly affecting children, adolescents, and young adults. Despite intensive multimodal therapy, metastatic EwS still remains associated with dismal outcomes (typically <30% overall survival rates) ^1,2^. Genetically, EwS is a rather ‘simple’ tumor driven by a single oncogenic driver, formed through fusing a member of the FET family of genes (*FUS*, *EWSR1*, *TAF15*) with an E26 erythroblast specific (ETS) transcription factor – in 85% of cases *FLI1*, resulting in the EWSR1::FLI1 fusion oncoprotein ^3–5^.

Accumulating evidence indicates that EwS cells reside in a dynamic metastable state characterized by fluctuating FET::ETS expression levels ^6–9^. Indeed, high *EWSR1::FLI1* expression is associated with a proliferative, epithelial-like phenotype, whereas low *EWSR1::FLI1* expression levels result in a more mesenchymal phenotype, accompanied by enhanced cell-matrix interactions, and increased migratory and metastatic capacity ^6,10–12^. These fluctuations have been linked to epithelial-mesenchymal plasticity ^12^, but the spatial architecture of heterogeneous fusion states within patient tumors and the downstream effectors promoting the EWSR1::FLI1-low metastatic phenotype remain elusive.

Spatial transcriptomic analyses in other cancer types reported that cancer cells at the leading edge of the tumor are more invasive than cancer cells inside the tumor core ^13,14^. In addition, genes associated with worse outcomes for cancer patients are part of the signature expressed at the invasive front in contrast to the tumor core ^15^. While single-cell and spatial transcriptomic studies in EwS are still scarce, a recent study employing spatial transcriptomics to investigate the tumor microenvironment (TME) has shed first light on the tumor-stroma crosstalk characterizing EwS tumors in the context of immune recruitment and proinflammatory microenvironmental signals ^16^.

In the present study, we show that the EWSR1::FLI1 activity in EwS is spatially heterogeneous, with high fusion-oncoprotein activity defining the tumor core and reduced activity predominating at the invasive tumor front. Spatial transcriptomics profiling revealed distinct transcriptional architectures between these regions, each associated with fusion-state dependent programs and clinical impact. Among the genes enriched within the EWSR1::FLI1-low signature (invasive front), we identified *LIM domain only protein 7* (*LMO7*) as a key downstream effector associated with poor overall patient survival and metastatic progression in a large EwS patient cohort. Integrated multi-omics and functional studies further establish LMO7 as a central regulator of epithelial-to-mesenchymal transition (EMT) and actin cytoskeleton remodeling, associated with EwS cell motility and metastatic behavior. Conditional silencing of *LMO7* decreased clonogenic growth and migration *in vitro*, and impaired tumorigenesis and metastasis *in vivo*, supporting a role of LMO7 as a clinically relevant mediator of oncogene-driven metastasis in EwS.

## RESULTS

### Spatially resolved tumor analysis reveals a decrease in EWSR1::ETS activity signature and high *LMO7* expression at the invasive front linked to metastasis in EwS

Metastasis is a complex multi-step process ^17^. In EwS, metastasis has been linked to a reduced activity of the EWSR1::FLI1 fusion oncoprotein ^6,12^. To identify key EWSR1::FLI1 downstream effectors potentially mediating metastasis programs *in situ*, we established an analytical pipeline integrating three transcriptomic datasets (**Fig. 1a**). The first dataset was derived from our previously published Ewing Sarcoma Cell Line Atlas (ESCLA), and comprised of five representative EwS cell lines: A-673, MHH-ES-1, SK-N-MC, and TC-71 harboring EWSR1::FLI1 fusions, as well as TC-106 driven by EWSR1::ERG ^18^. These cell lines were genetically modified to conditionally express a doxycycline (Dox)-inducible shRNA targeting the respective fusions allowing their silencing to around 20% remaining expression^18^. The second bulk-level transcriptomic dataset consisted of 196 clinically annotated EwS patient tumors with available overall survival data ^19^. FET::ETS silencing in these EwS cell lines induced strong and consistent upregulation (log_2_ FC >2.5) of 13 genes that were significantly associated with patient outcome (cut-off for each gene at the best percentile; log-rank *P* < 0.05, **Suppl. Table 1**). Focusing specifically on genes which high expression levels associated with poor prognosis yielded six candidates: *BMP1, CYR61*, *LMO7, LOX*, *NID2*, *PI15* (**Fig. 1b**). It should be noted that *LOX* had already been described as an EWSR1::FLI1-regulated tumor suppressor in the context of EwS ^20–22^. To refine this list, we reasoned that genes promoting EwS aggressiveness, and as such local tumor invasiveness, would display prominent spatial expression at the invasive tumor front, a region associated with elevated metastatic potential ^15^. Thus, we performed spatial transcriptomic analysis using the 10X Genomics Visium CytAssist Spatial Gene Expression platform on a localized gluteal EwS tumor (tumor purity ≈90%) from a 28-year old female patient (**Fig. 1c**). Spatial transcriptomic profiles were manually aligned to the CytAssist image and processed using a scanpy-based pipeline, enabling clustering based on transcriptional similarity while preserving the tissue architecture. This analysis resolved nine transcriptionally distinct clusters defined by their top marker genes, which faithfully recapitulated anatomical features such as tumor-adjacent muscle tissue (*DES, TTN*) and tumor-intrinsic, EWSR1::FLI1-regulated genes (*VEGFA, RBM11*) ^23,24^, thereby capturing the spatially organized anatomical and biological context (**Fig. 1d**). Tissue annotations were defined by overlaying hematoxylin and eosin (H&E) staining, clustering guided by spatial transcript expression (**Fig. 1d**), and inferred fusion transcriptional activity (**Fig. 1e**). Specifically, to define potentially invasive and more metastatic cells within the tumor mass, we inferred an established transcriptional signature of FET::ETS activity ^18^ and observed its strong reduction at the tumor invasive front (**Fig. 1e**). Among the six candidate genes, five (*BMP1*, *LMO7*, *LOX*, *NID2*, and *PI15*) were covered in the spatial transcriptomic panel and readily detected. Remarkably, LIM domain-only 7 (*LMO7*) exhibited the most prominent overexpression at the invasive front, inversely correlating with reduced FET::ETS fusion activity (*r* = –0.5254, *Padj* <0.0001) (**Suppl. Table 2**, **Fig. 1f, Suppl. Fig. 1a**). We then expanded and validated these results using a recently published cohort of 16 spatially profiled primary and treatment-naïve EwS tumors ^16^. Here, since zero-inflation for *LMO7* expression was remarkably higher than in our dataset, sections with fewer than 10 *LMO7*-expressing (non-zero) tumor spots were excluded from all subsequent analyses, in order to avoid artificial r/L inflation from underpowered sections (*n* = 4/16 excluded; *n* = 12/16 retained). Pooled analyses of these *n* = 12/16 samples confirmed the significant inverse association between *LMO7* expression and EWSR1::FLI1 transcriptional activity, evidenced by a negative correlation with the *EWSR1::FLI1*-high signature (*P*_EWSR1::FLI1high_ = 0.000043), and a positive correlation with the complementrary EWSR1::FLI1-low signature (*P* _EWSR1::FLI1low_ = 0.005) (**Suppl. Fig. 1b**). Interestingly, when these tumor samples were analyzed in detail for their *LMO7* expression distribution in non-zero tumor spots by evaluating whether neighborhoods that are locally core/intermediate/periphery-dense also tend to be locally *LMO7*-elevated, we observed a significantly higher Lee’s L at the periphery as compared to the intermediate regions in the section and especially the tumor cores, suggesting a preferential overexpression of *LMO7* at the invasive front of EwS tumors (**Fig. 1g**, Stouffer Z = –3.6, *P* = 0.000314, *n* = 11/16, one additional sample out of 12 was excluded from this comparison as it lacked *LMO7*-expressing spots in one of the two regions of interes, core and periphery).

**Figure 1.**
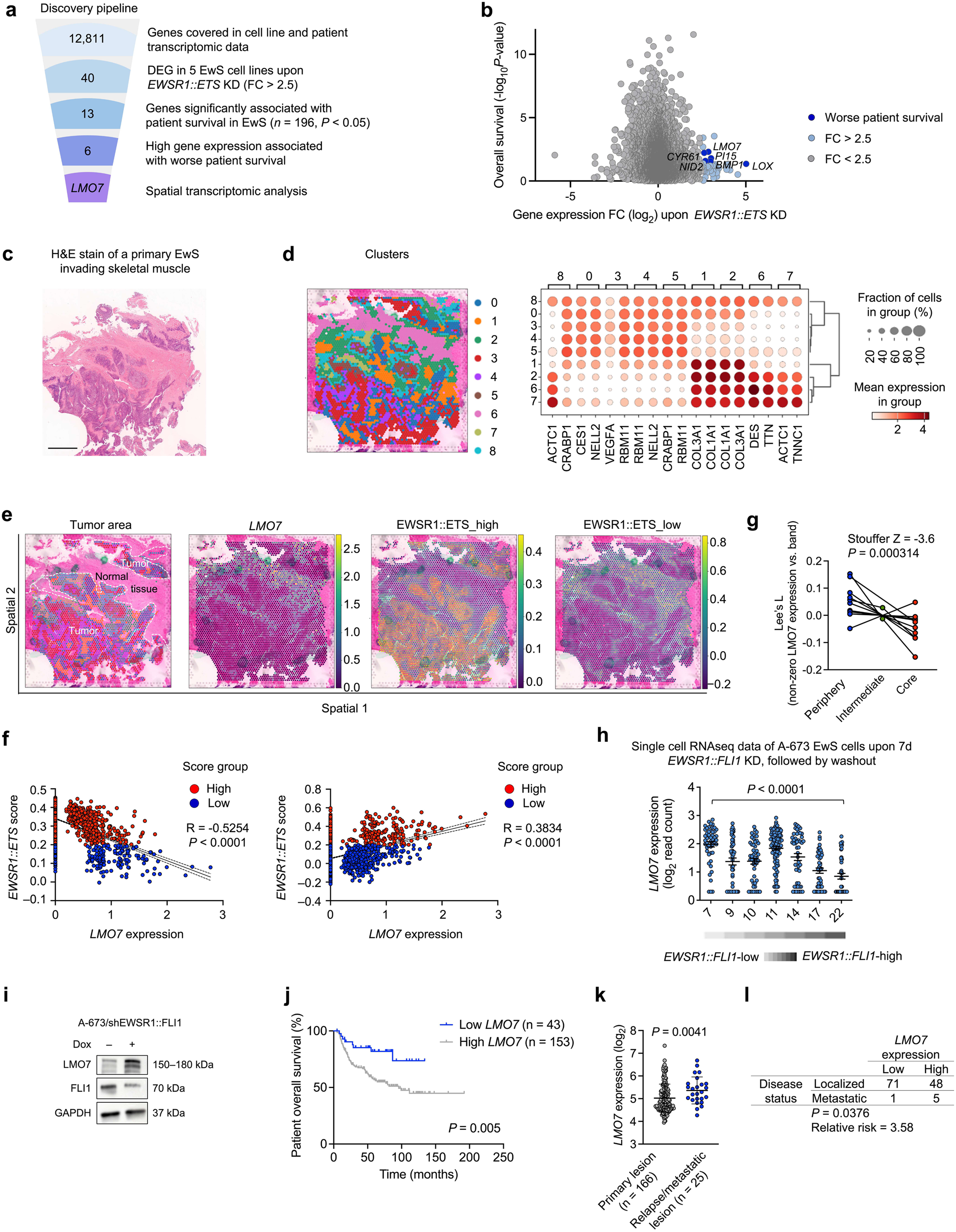
Spatially resolved analysis identifies enrichment of EWSR1::ETS-low activity signature and *LMO7* expression at the invasive tumor front linked to metastasis in EwS. **a.** Workflow depicting an integrative filtering approach to identify DEGs being highly expressed (FC > 2.5) in cell lines harboring a *EWSR1::ETS* KD, significantly associated with worse patient outcome, and highly expressed at the invasive front in an EwS patient sample profiled using spatial transcriptomics. Numbers represent the remaining gene candidates after each filtering step. **b.** Plot representing fold change (log_2_) expression of gene candidates in *EWSR1::ETS* KD cell lines (A-673, MHH-ES-1, SK-N-MC, TC-71, and TC-106) within the ESCLA dataset and their respective association with survival in an EwS patient cohort (n=196). Dark blue highlights six gene candidates (*BMP1*, *CYR61*, *LMO7*, *LOX*, *NID2* and *PI15*) displaying high expression in *EWSR1::ETS*-low ESCLA cell lines (FC > 2.5) and their expression being significantly associated with worse overall patient survival (*P* < 0.05). **c.** H&E staining of the EwS patient sample prior to spatial sequencing. **d.** Clustering of spatial data according to the top 10 marker genes in each spot. Circle size represents the fraction of cells in each cluster, the gradient of white-to-red color represents mean expression within the cluster. **e.** Spatial localization of clusters within the EwS patient sample. Tumor area was defined by the degree of transcriptional activity (median of gene expression counts) with minor adjustments according to the H&E staining in **c** and a EwS-specific signature ^18^). Second quadrant shows spatial expression of *LMO7*. Last two quadrants depict inferred spatial distribution of EWSR1::ETS activity signature from ^18^, according to the genes upregulated (*EWSR1::ETS*_high) or downregulated (*EWSR1::ETS*_low) by *EWSR1::ETS*. **f.** Spatial correlation of tumor-specific *LMO7* expression and EWSR1::ETS-high activity within defined tumor borders between (left, *r* = –0.5254, *P* < 0.0001), and *EWSR1::ETS*-low activity (right, *r* = 0.3834, *P* < 0.0001). Red dots depict cells with *EWSR1::ETS*-high, and blue depict cells with *EWSR1::ETS*-low activity signature. **g.** Slope plot comparing the median bivariate metric Lees’L of *LMO7* expression in non-zero tumor spots in *n* =11 spatially resolved EwS samples from ^16^ published dataset according to periphery, intermediate area, and tumor core regions. Stouffer Z and *P* value are depicted in the figure. **h.** Single cell expression of *LMO7* (RNA-sequencing) in A-673 EwS cell line with a conditional Dox-induced KD of *EWSR1::FLI1* upon Dox treatment for 7 d, followed by Dox washout in the next 15 d (GSE130025). White-to-black gradient indicates *EWSR1::FLI1* expression levels. Two-sided Mann-Whitney test between 7 d and 22 d, *P* < 0.0001. **i.** Western blot analysis of A-673 EwS cell lines upon Dox-mediated *EWSR1::FLI1* KD for 72 h. Antibodies against FLI1 and LMO7. Loading control: GAPDH. **j.** Kaplan-Meier survival analysis of 196 EwS patients stratified by best percentile of *LMO7* mRNA expression (Low, High). Two-sided Mantel-Cox test, *P* = 0.005. **k.** Relative *LMO7* expression of EwS primary tumors versus relapsed/metastatic lesions (*n* =191). **l.** Chi-square analysis of *n* = 125 EwS patients (exclusively primary lesions in patients with localized disease status, or metastatic lesions in patients with metastatic disease at diagnosis) dichotomized by median *LMO7* expression (low, high) (*P* = 0.0376).

In a next step, we further complemented these Visium-derived spatial data analyses, which does not fully allow for single-cell resolution, with a patient-derived single cell RNA-sequencing dataset from the Open Single-cell Pediatric Cancer Atlas (OpenScPCA) ^25^, restricted to four of seven patient samples with sufficient representation of both *EWSR1::ETS* tumor substates (≥10 cells per group; 17,874 cells total). Accounting for inter-patient variability with a mixed-effects logistic regression, *LMO7* detection was substantially higher in *EWSR1::FLI1*-low than *EWSR1::FLI1*-high tumor cells (OR = 18.3, 95% CI 6.47–51.6, P = 4.1 × 10⁻⁸; **Suppl. Fig. 1c**). In line with these data, re-analysis of a published single-cell RNA-sequencing dataset from A-673/shEF EwS cells with a Dox-inducible shRNA against *EWSR1::FLI1* ^26^ demonstrated a strong and significant upregulation of *LMO7* upon *EWSR1::FLI1* silencing, and *LMO7* expression inhibition upon *EWSR1::FLI1* reactivation (*P* < 0.0001) (**Fig. 1h**). Concordant results were obtained at mRNA and protein levels in bulk analyses (**Fig. 1i, Suppl. Fig. 1d**).

While the role of *LMO7* in EwS was unknown, it has been reported to regulate muscle differentiation genes ^27^, TGF-β signaling ^28^, and that it contributes to progression and metastasis of pancreatic ^29^ and breast carcinoma ^30^.

Consistent with the spatial data, high *LMO7* expression in our patient cohort (*n* =196) was associated with significantly worse patient overall survival (*P* = 0.005, Mantel-Cox test) (**Fig. 1j**). As previously shown in a subset of this cohort ^31^, the findings on *LMO7* were paralleled by an association of poor outcome in tumors with low fusion-activity scores in the new extended patient cohort (*n* = 196, *P* = 0.0006, Mantel-Cox test) (**Suppl. Fig. 1e,f**). Further, in this extended cohort, relapsed or metastatic lesions displayed a significantly higher *LMO7* expression level than primary tumors (*P* = 0.0041) (**Fig. 1k**). In addition, high *LMO7* expression was significantly associated with metastatic lesions in patients presenting with metastatic disease at diagnosis as compared with primary tumors from patients presenting with localized disease (*P* = 0.0376) (**Fig. 1l)**, an association that was only present for this gene candidate (**Suppl. Fig. 1g**). Collectively, these findings identify *LMO7* as a prognostically relevant gene in EwS, being specifically overexpressed at the invasive front of EwS tumors and transcriptionally suppressed by EWSR1::ETS.

### LMO7 mediates a pro-metastatic transcriptional state involving EMT, cytoskeletal remodeling, and integrin signaling in EwS

As there were – to the best of our knowledge – no reports on the role of *LMO7* in EwS available, we carried out gene set enrichment analysis (GSEA) coupled with weighted gene correlation network analysis (WGCNA) on *LMO7*-correlated genes within our transcriptomic data of 196 EwS tumors to obtain first clues on the potential function of LMO7 in EwS. These analyses revealed pleiotropic enrichments of locomotion/adhesion-related signatures including extracellular matrix remodeling, actin, collagen and integrin binding, and actin-based migration/movement (**Fig. 2a**), processes central to EwS tumor cell motility and metastatic dissemination ^6,32–34^.

**Figure 2.**
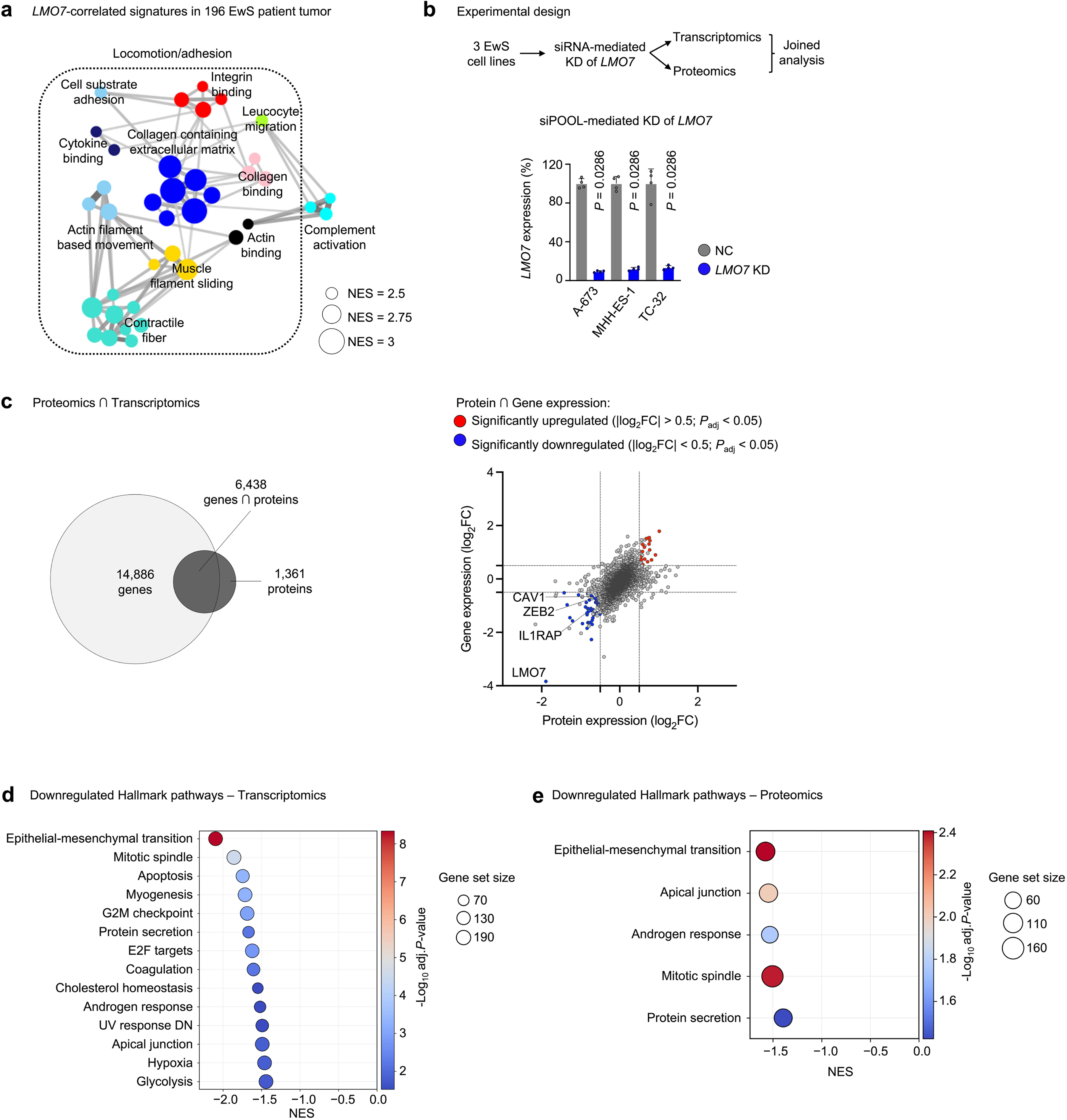
LMO7 defines a pro-metastatic transcriptional state involved in EMT and locomotion/adhesion signature enrichment. **a.** Weighted Gene Correlation Network Analysis (WGCNA) of enriched gene-sets obtained by Pearson correlation analysis of genes whose expression is positively correlated with *LMO7* expression in in Affymetrix expression data of 196 EwS tumors. Size of each node represents the Normalized Enrichment Score (NES), nodes belonging to the different pathways are represented with different colors. |NES| > 2, *P* < 0.05. **b.** Top: Experimental design for the matched transcriptomic and proteomic analyses upon *LMO7* siRNA-mediated KD. Bottom: *LMO7* mRNA expression in A-673, MHH-ES1 and TC-32 EwS cell lines upon siRNA-mediated LMO7 silencing, normalized to *RPLP0*. n = 4 biologically independent experiments. Horizontal bars represent the mean and whiskers the SD. Two-sided Mann-Whitney test. **c.** Left: Venn diagram depicting overlapping resulting genes and proteins. Right: Integrated analysis of the log_2_ foldchanges (log_2_FC) of commonly regulated genes/proteins derived from the transcriptomic and proteomic datasets. **d.** Gene set enrichment analysis (GSEA, Hallmarks gene set, MSigDB) of transcriptomic data derived from three EwS cell lines (A-673, MHH-ES-1 and TC-32) upon *LMO7* KD; |NES | > 1.5; *P*_adj_ < 0.05; Downregulated pathways. **e.** GSEA (Hallmarks gene set, MSigDB) of proteomic data derived from three EwS cell lines (A-673, MHH-ES-1 and TC-32) upon *LMO7* KD. |NES | > 1.5; *P*_adj_ < 0.05; Downregulated pathways.

To better discriminate which of these pleiotropic effects are directly mediated by *LMO7*, we performed matched transcriptomic and proteomic profiling of three EwS cell lines (A-673, MHH-ES-1, TC-32) following siPOOL-mediated *LMO7* knockdown (KD) or non-targeting negative control (NC) at a relatively early time point after transfection (55 h) to capture early, likely acute downstream changes rather than late potentially compensatory effects. As shown in **Fig. 2b**, the employed siPOOL led to less than 10–15% of *LMO7* residual expression, on average per cell line. Microarray-based transcriptome profiling (Affymetrix Clariom D arrays) captured 21,324 uniquely mapped transcript IDs, while the mass-spectrometry based proteomics yielded a set of 7,799 quantified proteins. Integrative analysis of these transcriptomic and proteomic datasets was performed to identify concordantly expressed genes/proteins. For this, only those genes/proteins represented in both omics’ modalities were retained (*n* = 6,438). A first analysis revealed that *LMO7* was the single most downregulated gene/protein across datasets (**Fig. 2c, Suppl. Fig. 2a,b)**, and that its downregulation led to the specific downregulation of other ECM-relevant transcripts including matrix metallopeptidase 2 (*MMP2*), synaptotagmin 11 (*SYT11*), and mitogen-activated protein kinase kinase kinase 12 (*MAP3K12*) (**Fig. 2c, Suppl. Fig. 2a,b, Suppl. Table 3,4**). In addition, the overlapping transcriptomic-proteomic output upon *LMO7* KD included interleukin 1 receptor accessory protein (IL1RAP), zinc finger E-box binding homeobox 2 (ZEB2), and caveolin-1 (CAV1) which had previously been described as mediators of metastasis in EwS ^35–37^, suggesting a regulatory interplay of *LMO7* with these targets in this context (**Fig. 2c**). Further, GSEA of pooled samples from the three tested EwS cell lines upon *LMO7* KD revealed EMT as the strongest downregulated pathway in Hallmark gene sets (MSigDB) at both transcriptomic and proteomic levels (**Fig. 2d,e**).

Collectively, these results indicate that LMO7 depletion results in downregulation of EMT signalling at mRNA and protein levels and suggests an involvement of LMO7 in the pro-metastatic signaling in EwS.

### Functional depletion of *LMO7* in EwS cells impairs clonogenic and spheroidal growth *in vitro* and tumorigenesis *in vivo*

To functionally investigate the role(s) of *LMO7* in EwS progression, we generated Dox-inducible shRNA expressing EwS cell lines (named hereafter A-673/shLMO7 and MHH-ES1/shLMO7) using two different shRNAs against *LMO7* (_1 and _2), and respective non-targeting controls. Dox-treatment led to efficient KD of LMO7 at the mRNA (**Fig. 3a**) and protein level (**Suppl. Fig. 3a**). Since clonogenic growth may reflect the outgrowth a of a single tumor cell to a colony similarly to the outgrowth of a single tumor cell to an overt metastasis, we first employed colony formation assays using the generated *LMO7* KD EwS cell lines. Here, *LMO7* KD resulted in a significant suppression of clonogenic growth (*P*_A-673/shLMO7_1_ = 0.0317; *P*_A-673/shLMO7_2_ = 0.0317; *P*_MHH-ES-1/shLMO7_1_ = 0.0286; *P*_MHH-ES-1/shLMO7_2_ = 0.0286; two-sided Mann-Whitney test) in *LMO7*-silenced EwS cell lines as compared to controls (**Fig. 3b**). To validate our results with independent approaches, we generated *LMO7* knockout (KO) models using Clustered Regularly Interspaced Short Palindromic Repeats (CRISPR) and *LMO7* overexpression models. Constitutive CRISPR-mediated KO of *LMO7* in A-673 cells resulted in similar levels of suppression of clonogenic growth (**Fig. 3c**). Conversely, conditional overexpression of *LMO7* led to an enhanced clonogenic capacity of A-673 cells (**Fig. 3d**).

**Figure 3.**
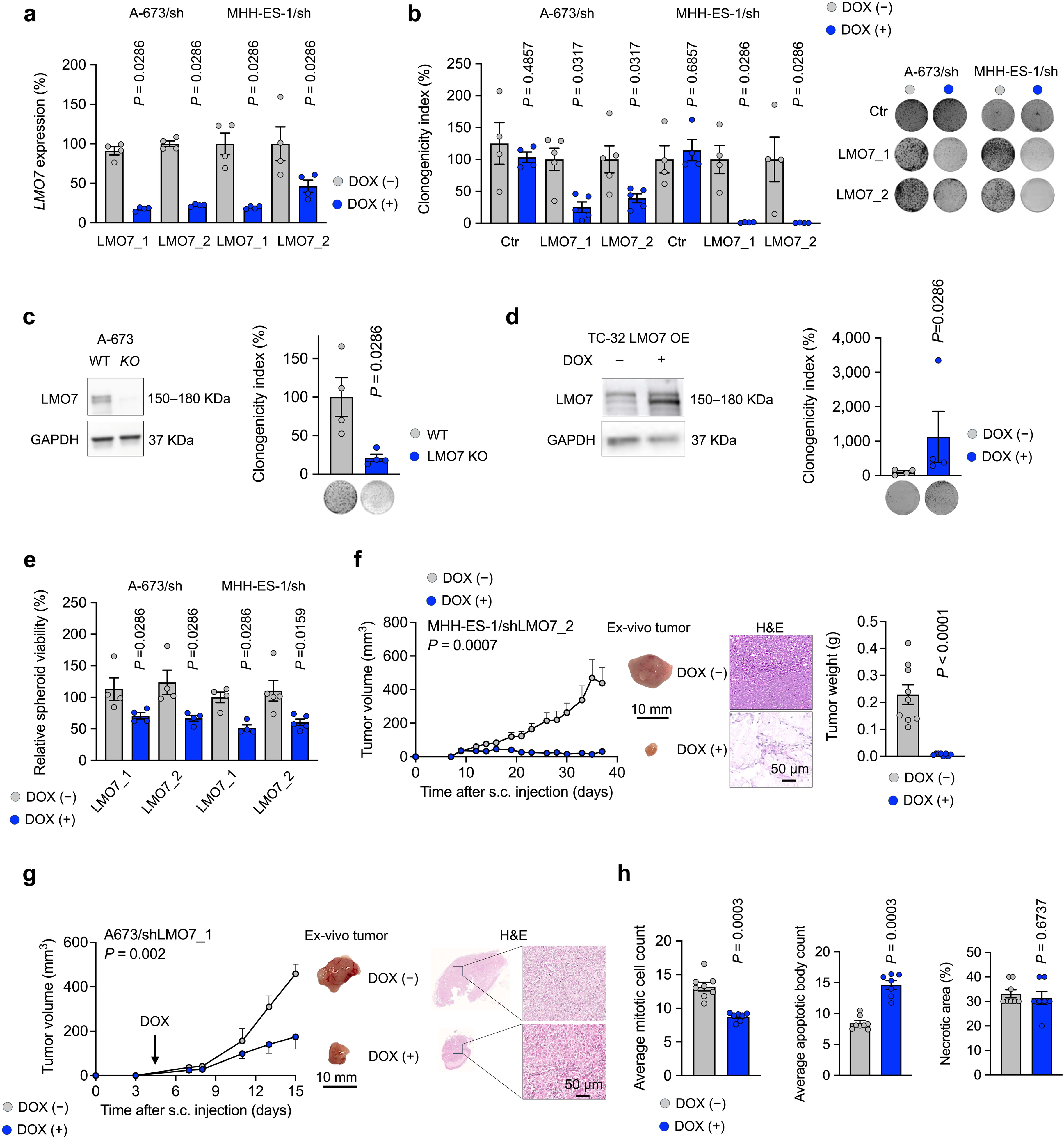
Functional depletion of *LMO7* in EwS impairs clonogenic growth and local tumor growth *in vivo*. **a.** *LMO7* mRNA expression upon Dox-inducible shRNA mediated LMO7 silencing (_1, _2) in A-673 and MHH-ES1 EwS cell lines, normalized to *RPLP0*. *n* = 4 biologically independent experiments. Horizontal bars represent the mean and whiskers the SD. Two-sided Mann-Whitney test. **b.** Left: Relative clonogenicity of A-673/shLMO7_1, A-673/shLMO7_2, MHH-ES-1/shLMO7_1 and MHH-ES-1/shLMO7_2 cells following 10 days of Dox-treatment (left) and representative colony images (right). *n* = 4 biologically independent experiments. Horizontal bars represent the mean and whiskers the SD. Two-sided Mann-Whitney test. Right: Representative images of clonogenic assays. **c.** Left: Representative western blot of LMO7 protein expression of constitutive *LMO7* KO in A-673 cell line (single cell clone). Right: Relative clonogenicity of A-673 wt and A-673-LMO7KO clone #22, following 10 days of Dox-treatment and representative colony images. *n* = 4 biologically independent experiments. Horizontal bars represent the mean and whiskers the SD. Two-sided Mann-Whitney test. **d.** Representative western blot of LMO7 protein expression in TC-32 EwS cells with Dox-inducible LMO7 OE (left). Relative clonogenicity of TC-32-LMO7 OE, following 10 days of Dox-treatment and representative colony images (right). *n* = 4 biologically independent experiments. Horizontal bars represent the mean and whiskers the SD. Two-sided Mann-Whitney test. **e.** Relative cell viability of A-673/shLMO7 (_1 and _2) and MHH-ES-1/shLMO7 (_1 and _2) spheroids, assessed by CellTiter-Glo assay, following 96 h treatment with/without Dox. *n* = 4 biologically independent experiments. Horizontal bars represent the mean and whiskers the SD. Two-sided Mann-Whitney test. **f.** Tumor volumes (mm^3^) upon subcutaneous injection of MHH-ES-1/shLMO7_2 with/without Dox-mediated *LMO7* silencing (left). *n* = 9 animals/condition. Horizontal bars represent the mean and whiskers the SD. Two-sided Mann-Whitney test for endpoint tumor sizes. Representative xenograft images and their respective H&E stains at 40× objective magnification (right). **g.** Measurements of tumor volumes (mm^3^) of subcutaneously injected A-673/shLMO7_1 with/without Dox-mediated LMO7 silencing (left). *n* = 8 animals/condition. Unpaired two-sided Mann-Whitney test for endpoint tumor sizes. *Ex vivo* representative images of xenografted tumors and respective hematoxylin-eosin stains. **h.** Average mitotic cell count, apoptotic bodies and necrotic area of tumors in **g**. *n* = 8 biologically independent xenografts/condition. Horizontal bars represent the mean and whiskers the SD. Two-sided Mann-Whitney test.

In line with these results, *LMO7* silencing reduced 3D sphere viability of A-673 and MHH-ES-1 EwS cells (**Fig. 3e**), which was mirrored by a significant delay of local tumor growth and weight of subcutaneous xenografts implanted in immunocompromised Nod/Scid/gamma (NSG) mice (*P*_A-673shLMO7_ = 0.002; *P*_MHH-ES1shLMO7_ = 0.0007; two-sided Mann Whitney test) (**Fig. 3f,g**). This reduction in tumor growth upon effective *LMO7* KD (**Suppl. Fig. 3b**) was accompanied in the immunohistochemically analyzed tumors by a significant reduction of mitotic index (*P* = 0.0003; two-sided Mann Whitney test), and a significant increase in the number of apoptotic bodies (*P* = 0.0003; two-sided Mann Whitney test), while no apparent differences in percentage of necrotic area were observed (**Fig. 3h**). In synopsis, these *in vitro* and *in vivo* results confirm a functional role of *LMO7* impacting tumorigenesis in EwS.

### *LMO7* silencing reduces EwS cell migration and metastatic burden in mice

Our previous omics-analyses of patient data and cell lines (**Figs. 2,3**) suggested a role of *LMO7* in migration and invasion. To test this possibility, we first carried out transwell migration assays of EwS cell lines with/without KD of *LMO7*. As shown in **Fig. 4a**, *LMO7* silencing significantly reduced cellular migration in two different cell line models (*P*_A-673shLMO7_1_ = 0.0286; *P*_A-673shLMO7_2_ = 0.0143 *P*_MHH-ES1shLMO7_1_ = 0.0143; *P*_MHH-ES1shLMO7_2_= 0.0143; one-sided Mann Whitney test). To further test the role of *LMO7* in metastatic spread *in vivo*, we employed a preclinical orthotopic xenograft mouse model of spontaneous metastasis formation ^38^. For this, we injected EwS cell lines harboring a conditional KD of *LMO7* into the tibia of NSG mice and monitored growth of primary tumors and metastases upon either Dox or control treatment (**Fig. 4b**). In this model, *LMO7* KD not only significantly suppressed primary tumor growth in the tibial bone (*P* = 0.0002, two-sided Mann-Whitney test; **Fig. 4b**), but also resulted in a significant inhibition of metastatic capacity *in vivo* in all animals tested as evidence by a significant decrease in liver weight (*P* = 0.0002; two-sided Mann-Whitney test) and the number of liver macrometastases confirmed by H&E stain (*P* = 0.0014; two-sided Mann-Whitney test) (**Fig. 4c**).

**Figure 4.**
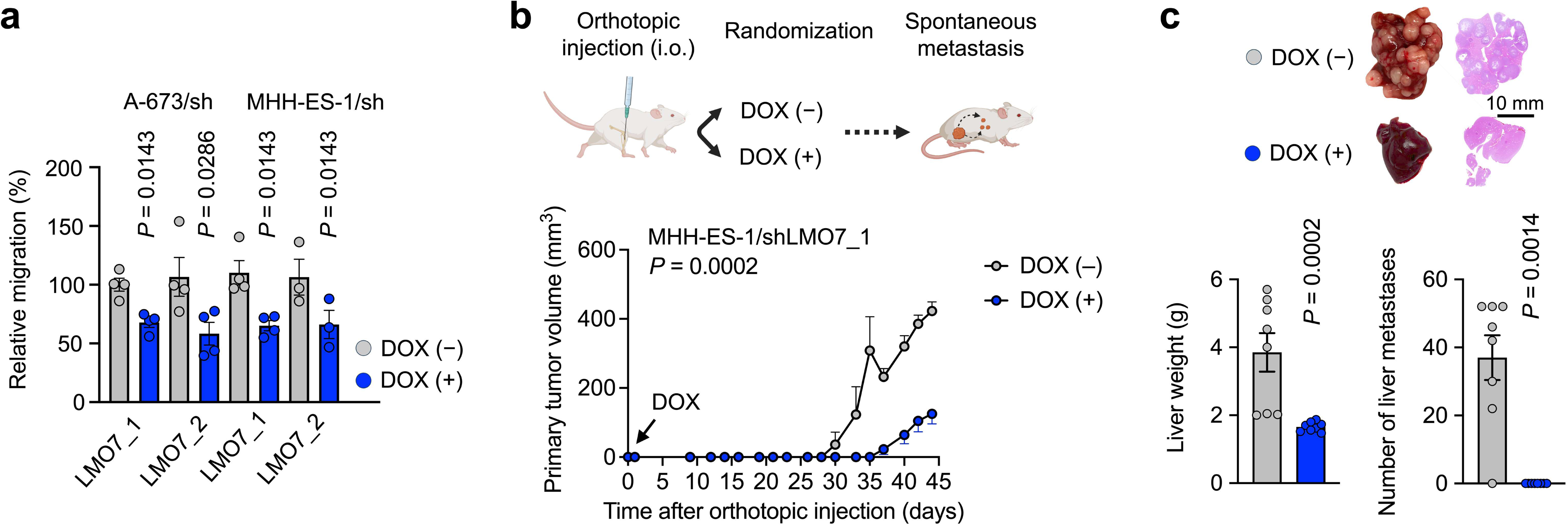
*LMO7* silencing reduces cell migration and metastatic burden in EwS. **a.** Relative cell migration of A-673/shLMO7 and MHH-ES-1/shLMO7 (_1 and _2) cells pretreated for 72 h with/without Dox. *n* ≥ 3 biologically independent experiments. Horizontal bars represent the mean and whiskers the SD. One-sided Mann-Whitney test. **b.** Top: Experimental design for an orthotopic mouse model of spontaneous metastatic spread. Bottom: measurements of leg volume (mm^3^) over time (days) in orthotopically injected MHH-ES-1/shLMO7_1 with/without Dox-mediated LMO7 silencing into right tibia (left). *n* = 8 biologically independent animals per condition. Unpaired two-sided Mann-Whitney test for endpoint tumor sizes. Created in BioRender. Aranaz, F. (2026) https://BioRender.com/249na7c. **c.** Top: representative images and IHC stains of mouse livers. Bottom: liver weight (g) of both groups at the experimental endpoint (left). Two-sided Mann-Whitney test. *n* = 8 animals/condition; number of hepatic macro-metastases observed in each mouse (right). Horizontal bars represent the mean and whiskers the SD. Two-sided Mann-Whitney test.

Collectively, these *in vitro*, *in vivo* and *in situ* results reveal *LMO7* as a spatially enriched effector downstream of EWSR1::ETS fusions at the invasive tumor front, that appears to orchestrate cytoskeletal dynamics, EMT, and metastatic progression in EwS.

## DISCUSSION

The biology of metastatic dissemination in EwS is only beginning to be systematically dissected and remains understudied relative to its primary tumor biology. A deeper understanding of the different cues that enable EwS cells to invade the surrounding tissue, and colonize distant organs is urgently needed, both to identify biomarkers for refined risk-stratification and to identify actionable liabilities in the metastatic cascade ^39^. In particular, there are a paucity of data on how EWSR1::ETS fusion activity is spatially modulated within the tumor microenvironment, and how such modulation might shape distinct cellular states with pro-metastatic properties.

A recent spatially-resolved study employing EwS primary material have pointed out at the presence of different transcriptional states shared by EwS tumor cells, including high *EWSR1::FLI1* fusion activity, and that these states colocalize with specific TME cells ^40^. In this study, we compliment these findings by providing additional evidence that the EWSR1::ETS activity signature is not uniformly distributed but instead exhibits a defined spatial organization within EwS tumors. Our results exhibit a zonal pattern, where a EWSR1::ETS-low transcriptional activity is enriched at the invasive tumor front, while the tumor core is characterized by a more pronounced EWSR1::ETS-high activity signature. This spatial segregation of fusion oncoprotein activity further supports the previous evidence that EwS cells dynamically tune their dependency on the EWSR1::ETS program in response to local microenvironmental cues, potentially zonally switching between proliferative, fusion-driven states, and more plastic, invasive states with attenuated EWSR1::ETS activity ^40^.

Strikingly, the EWSR1::ETS-low activity at the tumor leading edge is coupled to transcriptional induction of *LMO7*, in contrast to the tumor core, where high EWSR1::ETS activity coincides with comparatively low *LMO7* expression. This suggests that *LMO7* is embedded within a spatially restricted transcriptional program that favors invasion and dissemination of EwS cells. Given that LMO7 has been implicated in cytoskeletal organization, cell-cell adhesion, and mechano-transduction in other contexts ^41–43^, its upregulation at the invasive front of EwS tumors is consistent with a role in remodeling cell morphology and adhesion dynamics to facilitate tissue infiltration and metastatic spread. The observation that *LMO7* expression is tightly linked to a EWSR1::ETS-low state further supports the notion that that EwS cells transiently downregulate its oncogenic fusion activity to engage alternative transcriptional programs that support motility and extracellular matrix interaction ^6,7,9,12^.

Functionally, we demonstrate that LMO7 is a critical regulator of metastatic behavior in EwS *in vitro* and *in vivo*. Genetic and functional perturbation studies indicate that *LMO7* is necessary for migration and colonization, and its expression correlates with adverse clinical outcomes. High *LMO7* levels are associated with worse patient overall survival and a higher incidence of metastatic disease, underscoring its potential as a biomarker.

It has been reported that genes associated with worse outcomes for tumor patients are part of the signature expressed in the invasive front in comparison with the tumor core ^14,15^. Our results also support such a hypothesis and show that indeed in EwS, the invasive tumor front also expresses signatures that are associated with worse patient outcome. Zonal/regional expression of transcriptional programs suggests that EwS tumors may require at least a dual anticancer treatment for successful outcomes: one that would primarily target the invasive tumor front, and another that would mainly target the proliferative tumor core. Indeed, *in silico* modeling in oral squamous cell carcinoma showed that drug response predictions differ between tumor core and leading edge, suggesting that these regions may require distinct therapeutic strategies^15^.

Our results are in line with those from others describing the emerging role of *LMO7* as an important player in various cancers ^44^. For instance, *LMO7* was reported to promote progression and metastasis of breast ^30^ and pancreatic carcinoma ^29^, in the latter of which it has also been shown to drive immune evasion ^45^. Conversely, high *LMO7* expression was shown to correlate with improved outcome in oropharyngeal squamous cell carcinoma ^46^, and lung cancer ^47–49^, underscoring a context-dependent role of *LMO7*. Indeed, our data indicates the importance of *LMO7*-driven aggressiveness at the invasive front, demonstrating that high *LMO7* expression is associated with adverse prognosis and may serve as a novel candidate prognostic biomarker for negative outcomes in EwS. Hence, we propose to validate this finding in future studies in independent, prospective, clinical cohorts, to assess its robustness and potential utility as a prognostic marker.

More broadly, this work illustrates how integrating clinical data, spatially-resolved transcriptomics, multi-omic analyses, and functional models can uncover novel regulators of invasion and metastasis. Such approaches not only identify LMO7 as a clinically relevant effector in EwS but also provide a conceptual framework for discovering and prioritizing novel biomarkers which may offer inroads for targeted intervention.

Collectively, our results highlight *LMO7* as a central effector of a spatially defined, FET::ETS-low metastatic program in EwS, linking intratumoral zonation of fusion activity to clinically relevant metastatic potential.

## SUPPLEMENTARY FIGURE LEGENDS

**Supplementary Figure 1. Integrative multi-omics approach identified that EWSR1::ETS-low activity signature and *LMO7* expression are enriched at the invasive tumor front of EwS and are associated with worse patient outcome. a**. Spatial localization of clusters within the EwS patient sample. Tumor area (T) was defined by the degree of transcriptional activity (median of gene expression counts) with minor adjustments according to the H&E staining in **Figure 1c** and EwS-specific signature ^18^. Further images show spatial expression of candidate genes: *LOX*, *NID2*, *BMP1* and *PI15*. b. *LMO7* expression vs. *EWSR1::FLI1* activity (high or low) according to ^18^ in *n* = 12 spatially profiled EwS samples. Grey dots represent individual section *r* value, black bar is IQR, diamonds represent pooled median. Stouffer Z and *P* values are depicted for each analysis. **c.** Patient-derived single cell RNA-sequencing dataset from the Open Single-cell Pediatric Cancer Atlas (OpenScPCA) comprising four tumor samples. Cells were scored for high– or low-*EWSR1::FLI1* activity according to ^18^ (*P* = 4.13 ×10^-8^). **d.** Left: *EWSR1::FLI1* expression in EwS cells (A-673, MHH-ES-1 and SK-N-MC) containing a Dox-inducible construct for KD of *EWSR1::FLI1*, with/without treatment with Dox for 72 h. Right: *LMO7* expression upon KD of *EWSR1::FLI1* in same cell lines. n ≥ 4 biologically independent experiments. Two-sided Mann-Whitney test. **e.** Kaplan-Meier survival analysis of 196 EwS patients stratified by best percentile of inferred *EWSR1::FLI1* signature. Two-sided Mantel-Cox test, *P* = 0.0006. **f.** Kaplan-Meier survival analysis of 196 EwS patients stratified by best percentile of candidate gene (*LOX*, *NID2*, *BMP1*, *PI15* and *CYR61*) mRNA expression (Low, High). Two-sided Mantel-Cox test. **g.** Chi-square analysis of *n* = 125 EwS patients (exclusively primary lesions in patients with localized disease status, or metastatic lesions in patients with metastatic disease at diagnosis) dichotomized by median candidate gene (*LOX*, *NID2*, *BMP1*, *PI15* and *CYR61*) expression (low, high).

**Supplementary Figure 2. LMO7 defines a pro-metastatic transcriptional state involving in EMT and locomotion/adhesion signature enrichment. a.** Volcano plot showing the most significantly up– or down-regulated genes in transcriptomic analysis upon 55 h *LMO7* KD in three EwS cell lines (A-673, MHH-ES-1, TC-32). The blue dots indicate significantly differentially downregulated, while red upregulated genes (|log_2_FC | > 2; *P* < 0.05). **b.** Volcano plot showing the most significantly up– or down-regulated proteins in proteomic analysis upon 55 h *LMO7* KD in three EwS cell lines (A-673, MHH-ES-1, TC-32). The blue dots indicate significantly differentially downregulated, while red upregulated proteins (|log_2_FC | > 1; *P* < 0.05).

**Supplementary Figure 3. Functional depletion of *LMO7* in EwS impairs clonogenic growth and local tumor growth *in vivo*. a.** Representative western blot of remaining LMO7 protein expression upon Dox-inducible shRNA mediated LMO7 silencing using two independent shRNAs (_1, _2) in A-673 EwS cells. **b.** *LMO7* remaining expression in subcutaneous xenografts upon Dox treatment normalized to the housekeeping gene *RPLP0*. N = 8 tumors per condition. Horizontal bars represent the mean and whiskers the SD. Two-sided Mann-Whitney test

## DATA AVAILABILITY

The microarray and spatial sequencing data generated in this study are available via controlled access in the German Human Genome-Phenome Archive (GHGA, data.ghga.de) under the GHGA Accession code GHGAS62129574983531.

All proteomic data is deposited at the PRoteomics IDEntifications database (PRIDE) with accession code PXD080881. All other data supporting the findings of this study are available within the article and its supplementary information files.

## AUTHOR CONTRIBUTIONS

Veronika Buršić, Florencia Cidre-Aranaz, and Thomas G. P. Grünewald conceived the study. Veronika Buršić, Thomas G. P. Grünewald and Florencia Cidre-Aranaz wrote the paper and drafted all figures and tables. Veronika Buršić carried out *in vitro* experiments and statistical analyses. Veronika Buršić and Florencia Cidre-Aranaz performed *in vivo* experiments. Heng Luo, Clémence Henon, and Veronika Buršić performed spatial transcriptomic analyses. Lianghao Mao performed MS and raw data pre-processing. Clémence Henon, Angelina Yershova, Richard Arndt, Anna Ehlers, Veronika Buršić, and Florencia Cidre-Aranaz performed bioinformatics analysis. J-Ann M. Lego, Martha J. Carreno Gonzalez and Jing Li contributed to experimental procedures. Ana Sastre, Javier Alonso, Uta Dirksen, Moritz Gerstung, and Wolfgang Hartmann provided clinical samples or supervised bioinformatic analyses. Florencia Cidre-Aranaz and Thomas G. P. Grünewald supervised the study and data analysis. All authors read and approved the final manuscript.

## ACKNOWELEDGEMENTS

We thank Nadine Gmelin, Sabrina Knoth, Felina Zahnow, and Stefanie Kutschmann for their expert technical assistance. We thank Claudia Schmidt from the Preclinical and Translational Pathology Core Facility at DKFZ for conducting immunohistochemical stains. We thank Rainer Will from the Cellular Tools Core Facility at DKFZ for the assistance in generating the *LMO7* overexpression construct. We thank the DKFZ Microarray Core Facility for providing the Gene Expression Arrays and related services.

## ETHICS APPROVAL AND CONSENT TO PARTICIPATE

*In vivo* experiments were approved by the government of North Baden and conducted in accordance with ARRIVE guidelines and recommendations of the European Community (86/609/EEC) and UKCCCR (guidelines for the welfare and use of animals in cancer research). All tissue samples were retrieved from the biobank of the Cooperative Ewing Sarcoma Study (CESS). Analysis of human formalin-fixed, paraffin-embedded or cryopreserved tissue samples was approved by the ethics committee of the University of Heidelberg (approval no. S-211/2021).

## MATERIALS AND METHODS

### Provenience of cell lines and culture conditions

Human EwS cell lines were retrieved from the following responsories: A-673 (RRID: CVCL_0080) from American Type Culture Collection (ATCC), MHH-ES-1 (RRID:CVCL_1411) from the German Collection of Microorganisms and Cell Cultures (DSMZ), and TC-32 (RRID: CVCL_7151) from the Children’s Oncology Group (COG). Additionally, A-673, MHH-ES-1 and TC-32 cells modified with Dox-inducible construct targeting their FET::ETS fusion were previously generated as part of Ewing Sarcoma Cell Line Atlas (ESCLA) ^18^. Human HEK293T (RRID: CVCL_0063) cells were purchased from DSMZ. All cell lines were cultured in RPMI 1640 medium with stable L-glutamine and sodium bicarbonate (Sigma Aldrich, Germany), supplemented with 10% FCS tested to be doxycycline (Dox)-free (Sigma-Aldrich, Germany), penicillin (100 U/mL) and streptomycin (100 μg/mL; Merck, Darmstadt, Germany). Cell lines were routinely tested for Mycoplasma contamination using Mycoplasma PCR Detection Kit (Biozol, Hamburg, Germany) and checked for cell line identity and purity by short-tandem repeat (STR) or single nucleotide polymorphism (SNP) testing.

### Transient transfection

For transient silencing of *LMO7* expression in EwS cells, siPOOLs (siTOOLs Biotech GmbH, Planegg/Martinsried, Germany) were used consisting of 30 customized siRNAs against *LMO7* (si-G050-4008, siTOOLs Biotech GmbH), or non-targeting controls (si-C005). Cells were transfected with 5 nmol siPOOLs against *LMO7* or a non-targeting control siPOOL (both siTOOLs) using RNAiMax lipofectamine (Thermo Fisher Scientific, Darmstadt, Germany) in Opti-MEM (Thermo Fisher Scientific) as per manufacturer’s recommendation.

### Nucleic acid extraction and reverse transcription

Genomic DNA from human cell lines was extracted with the NucleoSpin Tissue kit (Macherey-Nagel, Düren, Germany), while plasmid DNA was extracted from bacteria ZymoPURE II™ Plasmid Midiprep Kit (Zymo Research Europe GmbH, Freiburg, Germany) as per manufacturer’s recommendation. Total RNA was isolated from EwS cell lines using RNA purification kit NucleoSpin RNA kit (Macherey-Nagel) by following the manufacturer’s protocol. RNA concentrations were measured on using Nanodrop One device (Thermo Fisher Scientific) and adjusted for 1 µg for reverse transcription using the High-Capacity cDNA Reverse Transcription Kit (Thermo Fisher Scientific) according to the manufacturer’s protocol for 20 µL reaction.

### Quantitative real-time polymerase chain reaction (qRT-PCR)

qRT-PCR reactions were performed using SYBR Select Master Mix for CFX (Applied Biosystems) mixed with 1:10 diluted cDNA, 0.5 µM forward and 0.5 µM reverse primer in a total reaction volume of 20 µl. The reaction was run on CFX Opus 96 Real-Time PCR System (Bio-Rad, Feldkirchen, Germany) and analyzed using Bio-Rad CFX Manager 3.1 software. Gene expression values were calculated using the 2^-(ΔΔCt)^ method relative to the housekeeping gene *RPLP0* as an internal control. Oligonucleotides were purchased from Sigma-Aldrich and are listed in **Suppl. Table 5**. The thermal conditions for qRT-PCR were as follows: UDG Activation (50 °C, 2 min) (1 cycle), AmpliTaq® Fast DNA Polymerase, UP Activation (95 °C, 10 min) (1 cycle); denaturation (95 °C, 15 sec) and annealing/extension (60 °C, 1 min) (45 cycles); denaturation (95 °C, 30 sec).

### Transcriptome analysis

RNA isolated from cell lines at 55 h post-transfection with siPOOLs (siTOOLs Biotech GmbH) silencing *LMO7* or non-targeting control and QC (> 50 ng/µl RNA per sample was available and the expression of *LMO7* and housekeeping gene *RPLP0* was tested using RT-qPCR) prior to submission to Microarray Core Facility (DKFZ, Heidelberg) for Affymetrix microarray analysis. The used Affymetrix human Clariom D (Applied Biosystems) captured a total of 135,750 transcripts. Firstly, *.cel files were processed using Transcriptome Analysis Console (Applied Biosystems), and output files were further processed using R studio/Bioconductor (R version 4.4.1). QC and further filtering showed 21,324 uniquely mapped transcript IDs with available Entrez IDs which excluded small, non-coding pseudogene or similar RNA transcripts. Principal Component Analysis (PCA) was applied to the transcriptomic count matrix after which principal components were subjected to Euclidean hierarchical clustering generating dendrograms representing relationships among sample groups. Differential expression analysis was performed on both pooled samples and cell line-specific groups using *limma* R package (3.62.2). Limma output files were ranked by differential gene expression and used as input for gene set enrichment analysis (GSEA). GSEA was performed using the clusterProfiler R package against the C2 gene set collection from MSigDB. Python (version 3.10.18, packaged by conda-forge) with Matplotlib and Seaborn libraries was used to visualize Hallmark gene set enrichment results in order to generate customized heatmap/bar plots representing enrichment scores and statistical significance.

### Generation of doxycycline (Dox)-inducible shRNA constructs

Human EwS cell lines A-673 and MHH-ES-1 were transduced with lentiviral Tet-pLKO-puro all-in-one vector system (plasmid #21915, Addgene) containing a puromycin-resistance cassette, and a tet-responsive element for Dox-inducible expression of shRNAs against *LMO7* (shLMO7_1 or shLMO7_2) or a non-targeting control shRNA (shCtr). shRNAs sequences are listed in **Suppl. Table 5**. Dox-inducible vectors were generated according to ^50^ and transformed into NEB® Stable Competent *E. coli* (High Efficiency) (New England Biolabs, Ipswich, MA, USA), and verified by sanger sequencing (**Suppl. Table 5)**. Lentiviral particles were generated in HEK293T cells. Virus-containing supernatant was collected to infect the human EwS cell lines. Successfully transduced cells were selected with 1 µg/ml puromycin (InvivoGen, San Diego, CA, USA). The shRNA expression for *LMO7* KD or expression of a negative control shRNA in EwS cells was achieved by adding 1 µg/ml Dox. Generated cell lines were designated as A-673/shCtr, A-673/shLMO7_1, A-673/shLMO7_2, MHH-ES-1/shCtr, MHH-ES-1/shLMO7_1 and MHH-ES-1/shLMO7_2.

### Generation of conditional LMO7 overexpression constructs

GeneArt Prime Clone gene synthesis (Thermo Fisher) was utilized to synthesize the longest *LMO7* transcript variant 1, normally expressed in EwS cells. *LMO7* transcript variant 1 was cloned into the Dox-inducible plasmid system rwSMARTTRE3G-LMO7-mCMV-TETON3G-Puro. Successfully transduced EwS cells were selected with 1 µg/ml puromycin (InvivoGen). The overexpression of *LMO7* in EwS cells was achieved by adding 1 µg/ml Dox to the growth medium.

### CRISPR-Cas9 *LMO7* KO

Cells were electroporated with Alt-R HiFi Cas9 Nuclease V3 (IDT, #1081061) and a two-part guide RNA (tracrRNA + crRNA): Alt-R CRISPR-Cas9 tracrRNA (IDT, #1072534) and LMO7 Alt-R® CRISPR-Cas9 crRNA (Hs.Cas9.LMO7.1.AE, 2nM, IDT). Cells were prepared by following the Alt-R™ CRISPR-Cas9 system protocol (IDT) for delivery of ribonucleoprotein complexes into cells using Lonza Nucleofector System (Lonza) and applying the DS-137 electroporation program. After electroporation, cells were plated in RPMI-1640 medium supplemented with 20% FCS for 72 h, and afterwards seeded as single cell clones. Once grown, clones were Sanger sequenced at Microsynth AG (Balgach, Switzerland) in the region of *LMO7* gene (**Suppl. Table 5**) and additionally confirmed for LMO7 protein presence/absence by western blot.

### Colony forming assays (CFAs)

For CFAs, 8×10^4^ cells/well were seeded in triplicate wells of a 6-well plate. After 24 h, Dox (1 µg/mL) was added to induce RNA-interference mediated *LMO7* KD. When performing CFAs with *LMO7* KO cell lines, parental wt EwS cell lines were used as control and standard medium was refreshed every 48 h. At the experimental endpoint (7–10 d after seeding, depending on the cell line), medium was removed, and cells were washed with DPBS (1×) (Gibco) and air dried for 5 min. Colonies were fixed with 800 µL/well ice-cold methanol (100%, Sigma-Aldrich) for 10 min at –20 °C. After removing methanol colonies were left to dry until white (5–10 min) and were then stained with 800 µL/well crystal violet solution for 20 min. After washing with DPBS (1×) and drying, plates were scanned and analyzed using ImageJ software (ImageJ 1.54f, NIH, USA).

### Spheroidal growth assays

For 3D sphere viability assays, A-673/shLMO7 and MHH-ES-1/shLMO7 cells were seeded in quadruplicates at a density of 1 × 10^3^ cells/well in a 96-well U-bottom, cell-repellent surface culture plates (Greiner Bio-One, Frickenhausen, Germany) with and without Dox (1 µg/mL). Every 48 h half of the media was renewed, adding fresh medium with 2× Dox. After 5 d, representative images of spheres were taken with 5× objective on an Axio light microscope with camera (Zeiss, Jena, Germany). Subsequently, sphere viability was measured using CellTiter-Glo Luminescent Cell Viability Assay (CTG) reagent (Promega, Madison, WI, USA). For that purpose, spheres were transferred to opaque plates and incubated with CTG reagent (1:1 ratio) in the dark on a shaker for 30 min. Luminescence from the samples was recorded at 1 sec integration time using a GloMax Microplate reader (Promega). Sphere viability was calculated by subtracting the blank values and relative to control.

### Transwell migration assay

Trasnwell experiments were performed as described in ^51^. Briefly, A-673/shLMO7 and MHH-ES-1/shLMO7 cells were pretreated with/without Dox (1 µg/mL) for 72 h in standard culturing conditions. Cells were then split and kept in 1% FCS (starvation) medium for 24 h in conditions with/without Dox (1 µg/mL). The next day, cells were seeded at a density of 5 × 10^4^ cells/well in 1% FCS cell culture media with/without Dox in Thincert cell culture insert for 24-well plates (pore size 8 µm, Greiner) and inserted into wells containing 10% FCS-supplemented medium with/without Dox (1 µg/mL). 24 h after seeding, migrated cells were fixed in 4% formalin and stained with crystal violet (Sigma-Aldrich). During staining, non-migrated cells were mechanically removed from the top of the insert using cotton swabs. Migrated cells were then destained using 10% acetic acid in PBS and absorbance of the solvent was measured using a GloMax Microplate reader (Promega) at 600 nm. The ratio of cells migrating toward the chemoattractant was calculated relative to control.

### Western blot

EwS cells were cultured in standard culture conditions, either with/without Dox (1 µg/ml) or wt and LMO7 KO cells for 72 h. Whole cellular protein was extracted using RIPA buffer and protease/phosphatase inhibitor cocktail (Sigma-Aldrich). Protein concentration was measured using the bicinchoninic acid assay (BCA assay) with Pierce BCA Protein Assay Kit (Thermo Fisher Scientific) and Pierce Bovine Serum Albumin Standards (Thermo Fisher Scientific), adjusted to 25 μg, separated on a 10% gel (10% Mini-PROTEAN® TGX™ Precast Protein Gels, 10-well – BioRad), and blotted on PVDF membrane using Trans-Blot Turbo Transfer System (BioRad). Membranes were blocked in 5% skim milk in 1 × TBST and incubated with rabbit polyclonal anti-LMO7 antibody (1:1,000, #PA5-5428, Invitrogen, Thermo Fisher Scientific), rabbit monoclonal anti-FLI1 (1:500, ab133485, abcam, Cambridge, UK), or rabbit monoclonal antiGAPDH (1:2,000, #14C10, Cell Signaling Technology (CST), Danvers, MA, USA), overnight at 4 °C. The next day, membranes were incubated for 1 h with mouse anti-rabbit IgG-HRP (1:5,000, #sc-2357, Santa Cruz Biotechnology, Dallas, TX, USA) in 5% skim milk at RT. Proteins signal was detected using WesternBright™ Sirius™ Chemiluminescent HRP Substrate (Advansta Inc., Menlo Park, CA, USA) and visualized with imaging system Fusion FX Edge (Vilber Lourmat, Marne-la-Vallée, France) equipped with Evolution CAPT Edge software.

### Mass spectrometry and global proteome analysis

Cells were seeded at 1.4 × 10^5^ cells/well in 6-well plates in four biological replicates for global proteome analysis. The next day, cells were incubated with siPOOL targeting either *LMO7* or non-targeting control. At 55 h post-transfection, cells were washed twice with cold DPBS (1×), the plates transferred to ice, and cells lysed with 100 µL 1% sodium deoxycholate-based buffer per well. Cells lysates were transferred to 1.5 mL locked Eppendorf tubes using a cell scraper and boiled at 100 °C, 10 min. Samples were then cooled on ice and stored at –80 °C until processing as described in ^52^. In brief, heat-denatured and sonicated proteins were digested for 16 h with trypsin and LysC, and peptides were purified using Styroldivinylbenzol-Reversed Phase Sulfonat (SDB-RPS) stage tips, then dried and reconstituted for quantification. For mass spectrometry, 400 ng of peptides were separated on a nanoElute system (Bruker Daltonics Inc, Bremen, Germany) coupled with a TIMSTOF HT mass spectrometer (Bruker Daltonics, Bremen, Germany) operating in DDA-PASEF mode with a 120 min gradient. Data were processed using FragPipe (Version 20) with label-free quantification and match-between-runs enabled, referencing the UniProt human database with a 1% FDR threshold.

Downstream bioinformatics analyses were conducted in Perseus (version 1.6.7.0) and in R (version 4.4.2). In R, raw MaxLFQ intensities were imported and processed with the DEP Bioconductor package (version 1.28.0) following the workflow of Zhang et al. (2018). Protein intensities were normalized using variance-stabilizing normalization (version 3.74.0) to remove systematic bias. Missing values (assumed to be “missing-not-at-random”) were imputed by sampling from a left-shifted Gaussian distribution via DEP’s MinProb algorithm (q = 0.01). Differential expression between LMO7 KD and NC was carried out with the *limma* package (version 3.62.2). Linear models were fitted (lmFit), empirical Bayes moderation applied (eBayes(…, robust = TRUE)), and the KD-NC contrast extracted (makeContrasts, contrasts.fit, eBayes). Data manipulation and table joins (e.g. matching gene IDs) were performed using the tidyverse backend (dbplyr v1.1.4), and gene annotations sourced from org.Hs.eg.db (version 3.20.0).

Ranked log₂ fold changes were then subjected to GSEA using Hallmark gene sets obtained via MsigDB (version 25.1.1). Pathways with FDR adjusted *P*< 0.05 were considered significant, and the top 10 hallmark pathways by absolute normalized enrichment score (NES) were visualized using Python (version 3.10.18, packaged by conda-forge) with Matplotlib and Seaborn libraries in order to generate customized heatmap/bar plots representing enrichment scores and statistical significance.

### *In vivo* experiments

NOD/Scid/gamma (NSG) mice were maintained in individually ventilated cages (IVC) under specific pathogen-free (SPF) conditions with strict dark/light cycles (darkness from 8 p.m. to 6 a.m.), an ambient temperature of 20–24 °C and a humidity of 45–65%. Subcutaneous xenograft experiments were performed as described in ^53^. Briefly, 2 × 10^6^ EwS cells per animal were suspended in 1:1 DPBS (1×):Geltrex ratio and injected in the right flank of 10–12 weeks old NSG mice. Both tumor diameters were measured every second day with a caliper, and tumor volume was calculated by the formula (L × l^2^) / 2, where L is the length and l the width. When the tumors reached an average volume of ∼100 mm^3^, mice were randomly distributed in equal groups. The control group received 5% sucrose (Sigma-Aldrich) in drinking water, while the treatment group received 2 mg/mL Dox BelaDox (Bela-pharm) with 5% sucrose (Sigma-Aldrich). Before tumors in the either group reached the volume of 15 mm in either direction, all mice were killed by cervical dislocation. Other humane endpoints were determined as follows: Ulcerated tumors, loss of ≥20% body weight, constant curved or crouched body posture, bloody diarrhea or rectal prolapse, abnormal breathing, severe dehydration, visible abdominal distention, obese Body Condition Scores (BCS), apathy, and self-isolation.

For orthotopic xenograft experiments, 24 h before injection mice were treated with 800 mg/kg mouse weight/day Metamizole in drinking water as analgesia. On the day of injection, mice were anesthetized with inhalable isoflurane (1.5–2.5% in volume) and their eyes were protected with Bepanthen eye cream. After disinfection of the injection site, 2×10^5^ EwS cells harboring a shRNA against *LMO7* in 20µl DPBS (1×) were injected using a 30 G needle (Hamilton, USA) into the right proximal tibia. Mice were subsequently treated with Metamizole in drinking water (800 mg/kg mouse weight/day) for 24h. At that point, mice were randomized in equal groups and either received 5% sucrose (Sigma-Aldrich) in the drinking water (controls) or 2 mg/ml Dox BelaDox (Bela-pharm) with 5% sucrose (treatment group). Primary tumor growth was followed by measuring the leg circumference every second day using a caliper, and tumor volume was calculated with the formula (L × l^2^) / 2 as before. Animals were killed by cervical dislocation at the predefined experimental endpoint described previously, or if they reached a humane endpoint as listed above or exhibited signs of limping at the injected leg (event). After extraction of the tumors, a small fraction of each tumor was snap frozen in liquid nitrogen to preserve the RNA isolation, while the remaining tumor tissue was fixed in 4%-formalin and embedded in paraffin for immunohistology. In the case of the orthotopic model, all inner organs were harvested, weighed, photographed, 4%-formalin-fixed and embedded in paraffin for (immuno)histology. For analysis of the extent of metastatic spread, HE-stained histological slides were evaluated for presence of EwS cells.

### Immunohistochemistry (IHC)

Formalin-fixed tumors and metastases were dehydration following standard protocols. Bone samples were subsequently decalcified using the Decalcifying Solution-Lite (Sigma Aldrich) for a week at 4 °C. Hematoxylin and eosin (H&E) stained sections were analyzed for the percentage of tumor necrosis over the total tumor area, as well as for mitotic and apoptotic index by counting 10 high-power fields (40×) per sample.

### EwS patient cohort analysis

Kaplan-Meier survival analyses were performed on 196 EwS tumor samples that had been molecularly confirmed and retrospectively collected, as well as profiled at the mRNA level by gene expression microarrays in previous studies ^54–57^. Microarray data was generated using Affymetrix HG-U133Plus2.0, Affymetrix HuEx-1.0-st or Amersham/GE Healthcare CodeLink microarrays and is associated with following Gene Expression Omnibus (GEO) accession codes: GSE34620 ^54^, GSE17618 ^57^, GSE12102 ^56^, GSE63157 ^55^, provided with clinical annotations data was normalized separately as previously described ^58^. Genes represented on all microarray platforms were kept for further analysis. Batch effects were removed using the ComBat algorithm ^59^ and data processing was done in R. For Kaplan-Meier analyses of overall survival from 196 EwS patient cohort, statistical differences between the groups were assessed by a Mantel-Cox test.

### Gene set enrichment analysis (GSEA)

To identify enriched gene-sets associated with LMO7 gene expression in primary tumors, we created a pre-ranked list of genes ordered by Pearsońs correlation coefficient with LMO7 based on transcriptomic data of 196 EwS. GSEA was performed in R (version 4.3.0) using the C5 ontology gene sets from MSigDB and using the pre-ranked list as input. In order to construct a network, the Weighted Gene Correlation Network Analysis R package (WGCNA R) ^60^ was used as described in ^61^. Briefly, a binary matrix of GO-terms × genes (where 1 indicates the gene is present in the GO term and 0 indicates it is not) was created. Then, the Jaccard’s distance for all possible pairs was computed to create a symmetric GO adjacent matrix. Clusters of similar GO terms were identified using dynamicTreeCut algorithm, and the top 20% highest edges were selected for visualization. The highest scoring node in each cluster was determined as the cluster label (rName). The obtained network and nodes files were fed into Cytoscape (v 3.10.0) for network design and visualization.

### Integrated transcriptomic and proteomic analysis

To identify concordantly expressed genes/proteins, only those expressed at both the mRNA and protein level were retained (*n* = 6,438). Expression distributions were examined via histograms and density plots, with Pearson’s correlation quantifying transcriptomic-proteomic log_2_ FC relationships (*r* = 0.594, *P* < 0.0001). A quadrant-based classification system categorized genes and proteins based on expression patterns using platform-specific thresholds (|log_2_FC| > 1 for genes, |log_2_FC| > 0.5 for proteins, adjusted *P* < 0.05 for both. Data processing and filtering was handled with dplyr function for identifying the variables to plot. Analysis was performed using RStudio (R 4.4.1).

### Spatial transcriptomics using 10X Visium CytAssist spatial gene expression platform

A 10 µm sample section was cut from a EwS FFPE sample (KITZ-SARC-0191) using Microm HM 355S and placed on a SuperFrost slide. The sample was dried overnight in a desiccator to eliminate residual moisture after which it was stained with H&E and destained following protocol CG000520 Rev C from 10X Genomics. After decrosslinking, the sample was processed using the 10X Genomics FFPE Spatial Gene Expression 6.5 mm Human Transcriptome Kit (Catalog No. 1000443) following the manufacturer’s protocol (CG000495 Rev F) with the Visium CytAssist platform. The final Visium sequencing library was sequenced on the NovaSeq™ X Series platform using the 1.5B Reagent Kit (100 Cycles) following the manufacturer’s protocol. The raw sequencing output was first converted into FASTQ files using the ‘mkfastq’ function in Space Ranger (version 3.0.0, 10X Genomics). The aligned FASTQ files were subsequently processed using the ‘count’ function in Space Ranger (version 3.0.0, 10X Genomics) to generate the gene expression count matrix, using the GRCh38-2020-A reference genome and the Visium Human Transcriptome Probe Set v2.0 (GRCh38-2020-A). The spatial transcriptomics count data was manually aligned with CytAssist image using the Loupe Browser.

Spatial transcriptomic data was analyzed using the scanpy workflow ^62^. Input data contained 4,503 observations (spatial location on the tissue) and 18,085 gene variables, n_obs × n_vars = 4,503 × 18,085. Quality control steps included the filtering of spatial features based on min and max count (min_counts = 1,000, max counts = 35,000) as well as filtering out those with >20% mitochondrial genes. Those steps yielded 4,362 spatial locations for downstream analysis. Genes that were detected in less than 10 spots were also filtered out (*n* = 84). After normalization and logarithmic transformation of the data, 5,000 highly variable genes were selected using scanpy.pp.highly_variable_genes() function with ‘seurat’ flavor. Dimensionality reduction was performed to summarize the data with 50 principal components, followed by the construction of a neighborhood graph to model local similarities. Clustering was conducted using the Leiden algorithm (with resolution = 1), and results were visualized in both UMAP and spatial coordinates overlaid on tissue images to preserve anatomical context. Data from analyzed tumor marker expression, high transcriptional activity and H&E staining was used to define the border of tumor and tumor-adjacent tissue in the sample. Then within defined tumorous tissue, spatial expression of genes of interest was correlated to the expression of genes know to be targets downregulated (EWSR1::ETS_low) or upregulated (EWSR1::ETS_high) by the EWSR1::ETS fusion, available from ^18^. Fusion activity signature was then overlaid on the tissue according to the spatial coordinates of the analyzed sample. The analysis was run in Python version 3.10.18 (packaged by conda-forge).

### Spatial transcriptomics analysis of published 10X Visium EwS dataset

Spatial transcriptomic data for 16 Ewing sarcoma sections were obtained from the publicly deposited Visium dataset ^16^, comprising raw Space Ranger count matrices and spot coordinates. Spots with fewer than 500 UMIs, fewer than 200 detected genes, or more than 30% mitochondrial reads were excluded; counts were normalized to 10,000 per spot and log-transformed. No gene-level filtering was applied, to preserve coverage of downstream signature genes. *LMO7* expression was detected in a median of 7.4% of tumor spots per section (mean 9.7%; range 1.3–34.3%), consistent with pronounced zero-inflation. All analyses therefore used *LMO7* expression from spots with detectable transcript only, treating non-expressing spots as missing data rather than true zero values. Sections with fewer than 10 *LMO7*-expressing tumor spots were considered too sparse for a stable estimate and were excluded from the primary pooled analysis (12 of 16 sections retained).

As a preliminary non-spatial analysis, *LMO7* expression was directly correlated (Pearson *r*) with the two *EWSR1::ETS_high* and *EWSR1::ETS*_low transcriptional activity signatures ^18^, each scored per spot with UCell; Pearson correlations were computed across nonzero-*LMO7* tumor spots within each section. Per-section significance was assessed by permutation (999 permutations of the signature score, *LMO7* values held fixed), yielding an empirical two-sided *P* value, and results were pooled across sections using Stouffer’s method (per-section *P* values converted to signed z-scores and combined into a meta-analytic z-statistic and *P* value).

For further correlation between *LMO7* expression and EwS spatial pattern, we performed an unbiased tumor core/intermediate/periphery region of interest classification. To this aim, we first classified spot clusters as tumor vs non-tumor. For this, all 16 sections were integrated: the 2,000 most variable genes were selected ^63^, expression was scaled and reduced by PCA (30 components), and batch effects were corrected with Harmony ^64^. Unsupervised clustering was performed with the Leiden algorithm ^65^ at resolution 0.4, matching the original study. Each cluster was classified as tumor or non-tumor based on its median expression of a six-gene *EWSR1::FLI1* tumor marker score (*NKX2-2*, *CD99*, *CAV1*, *CCND1*, *HES1*, *PAPPA*; scored per spot with UCell); clusters above a fixed threshold were designated tumor, and all remaining clusters (e.g., stromal, immune, and vascular spots) were pooled as non-tumor. Individual spots inherited the classification of their assigned cluster.

Next, to delineate tumor-periphery spatial regions, a spatial neighbor graph (six nearest neighbors on the hexagonal Visium lattice; Squidpy, ^66^ was constructed for each section independently. For each tumor spot, the shortest-path (graph-hop) distance to the nearest non-tumor spot was computed; tumor spots with no finite path to stromal tissue were excluded. Tumor spots were stratified into tertiles of this distance to define three regions: a peripheral band (nearest tertile to non-tumors regions), an intermediate band, and a core band (farthest tertile). Bands were thus defined by graph distance to annotated stromal spots, not by cell-type deconvolution.

Spatial co-localization between *LMO7* expression and each band was quantified using the bivariate spatial association statistic of Lee (2001), which jointly captures the correlation and spatial autocorrelation of two variables. For each comparison, a binary band-membership indicator and *LMO7* expression were independently standardized (zero mean, unit variance) across all in-tissue spots of the section, and the statistic was computed as

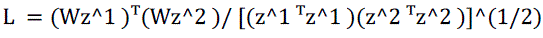

where z₁ and z₂ are the standardized variables and W is the row-normalized spatial weight matrix from the same neighbor graph. Statistical significance per section was assessed by permutation testing (999 permutations of *LMO7* expression, graph and band assignments held fixed), yielding an empirical two-sided p-value. Section-level results were pooled using Stouffer’s method: per-section *P* values were converted to signed z-scores and combined into a single meta-analytic z-statistic and *P* value. For the core-versus-periphery comparison specifically, one additional section was excluded because none of its *LMO7*-expressing spots fell within the core band, yielding a final set of 11 sections.

All analyses were performed in Python version 3.12 using Scanpy ^62^ and Harmony ^64^ for preprocessing and batch integration, Squidpy ^66^ for spatial graph construction, and SciPy and statsmodels for permutation testing and pooled statistics.

### Single-cell analyses of publicly availabled 10X scRNA-seq EwS dataset (ALSF)

Single-nucleus RNA-seq data were downloaded from the ScPCA Portal (Childhood Cancer Data Lab)^25^; project SCPCP000015 (accessed December 2025; 13 libraries, 10x Chromium v3.1, nuclei). Six libraries are O-PDX models ^67^ and were excluded a priori. Analysis was restricted to the seven patient-derived libraries: SCPCL000822, SCPCL000824, SCPCL000825, SCPCL000826, SCPCL000827, SCPCL000828, SCPCL001111. For each library, the ScPCA-provided filtered SingleCellExperiment object was converted to a Seurat object. Barcodes with openscpca_celltype_annotation == “openscpca-excluded” were removed. Counts were log-normalized with NormalizeData (default: total-count scaling to 10,000, natural-log of 1+x).

Tumor cell identity and *EWSR1::FLI1* substate were assigned upstream by OpenScPCA (cell-type-ewings module, ref; data release 2024-11-24) and used without re-annotation. Substate calls are AUCell-threshold-based: EWS-high requires AUC above threshold for both the Aynaud EWSR1::FLI1 direct-target gene set ^26^ and MSigDB STAEGE_EWING_FAMILY_TUMOR; EWS-low requires AUC above threshold for both the Wrenn NT5E-associated mesenchymal gene set and HALLMARK_EPITHELIAL_MESENCHYMAL_TRANSITION. Cells meeting neither or both criteria, and the EWS-high-proliferative subclass, were excluded from this comparison, leaving only EWS-high and EWS-low cells.

*LMO7* detection was defined per cell as log-normalized count > 0. Patients were included only if they contributed ≥10 EWSR1:ETS_high and ≥10 EWSR1::ETS_low cells; 4/7 patients met this criterion (SCPCL000822: 2,778 high / 639 low; SCPCL000824: 6,002 high / 93 low; SCPCL000825: 4,782 high / 21 low; SCPCL000826: 3,483 high / 76 low; 17,874 cells total). Detection was modeled by mixed-effects logistic regression, where patient was specified as a random intercept and random slope on EWSR1::ETS status to account for between-patient variation in both baseline detection and effect size.All analyses were performed with R version 4.5.1.

