## Supplementary Figure 1 for "Functional spatial transcriptomics uncover LMO7 as a fusion-regulated and clinically relevant driver of metastasis in Ewing sarcoma"

**Supplementary Figure 1**. Bursic *et al.*

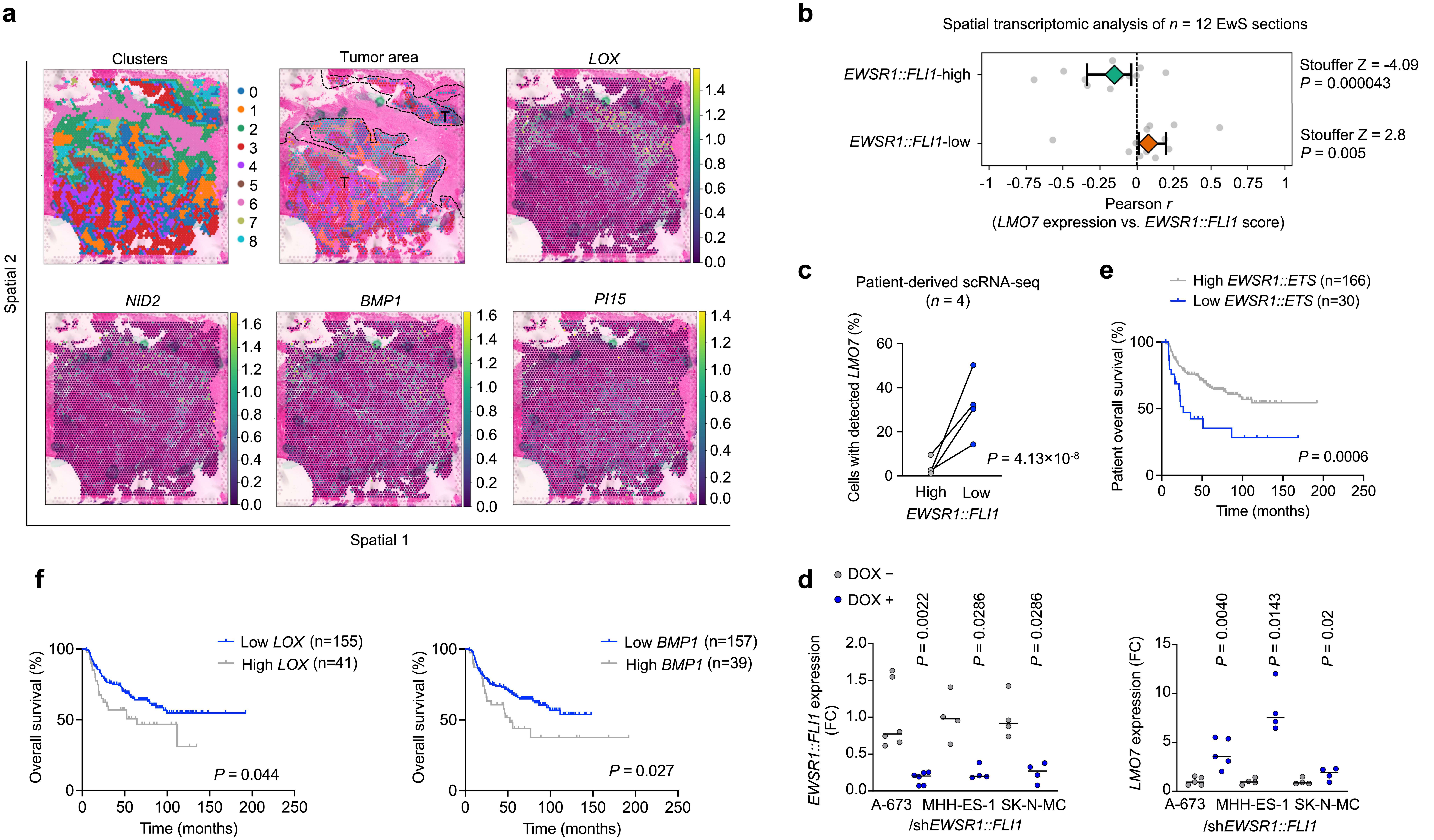


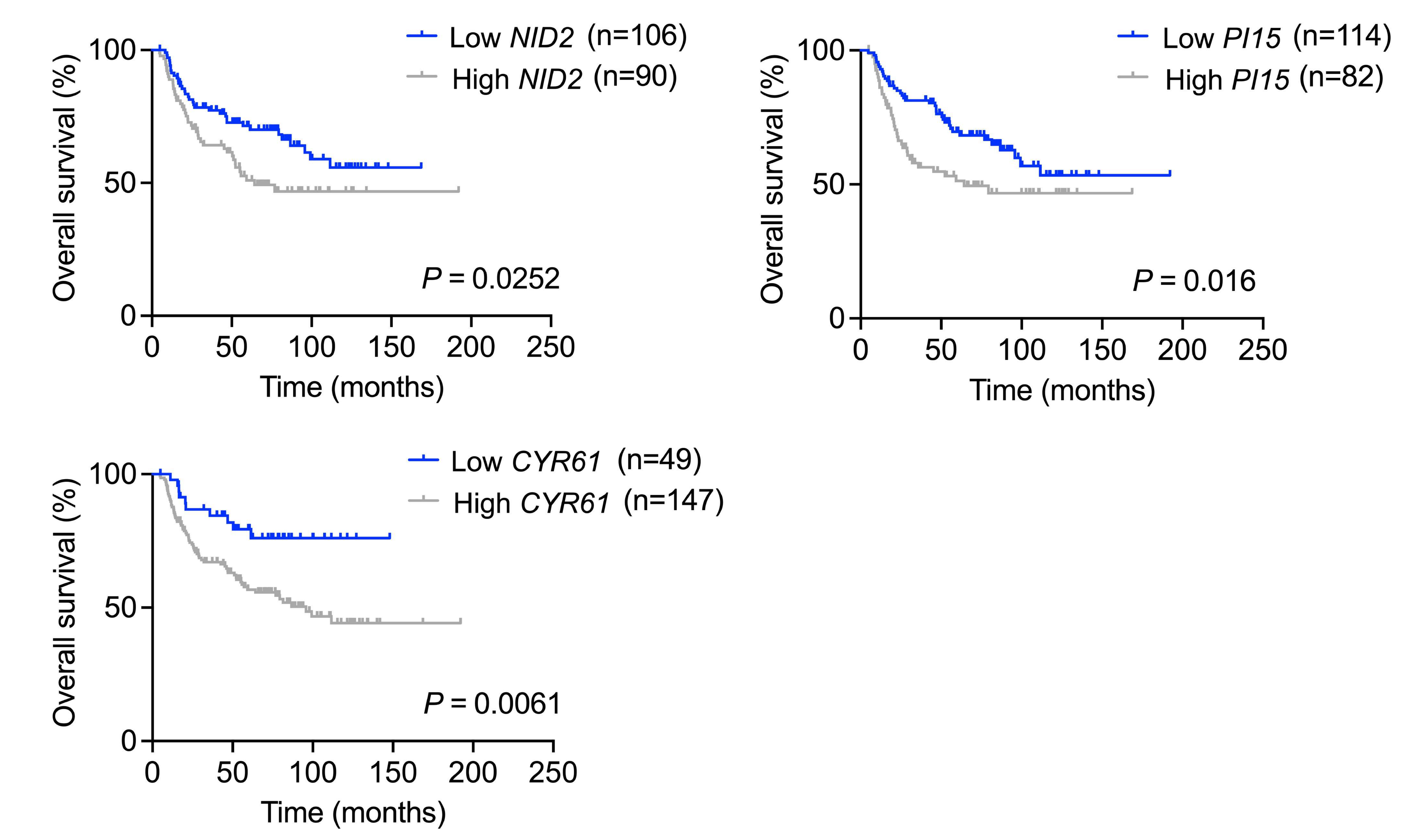


**g**

| | | *LOX* expression | |
|---|---|---|---|
| | | Low | High |
| Disease status | Localized | 63 | 56 |
| | Metastatic | 2 | 4 |

*P* = 0.8798
Relative risk = 1.588

| | | *NID2* expression | |
|---|---|---|---|
| | | Low | High |
| Disease status | Localized | 64 | 55 |
| | Metastatic | 2 | 4 |

*P* = 0.3276
Relative risk = 1.613

| | | *CYR61* expression | |
|---|---|---|---|
| | | Low | High |
| Disease status | Localized | 62 | 56 |
| | Metastatic | 4 | 2 |

*P* = 0.4988
Relative risk = 0.7881

| | | *PI15* expression | |
|---|---|---|---|
| | | Low | High |
| Disease status | Localized | 63 | 56 |
| | Metastatic | 3 | 3 |

*P* = 0.888
Relative risk = 1.059

| | | *BMP1* expression | |
|---|---|---|---|
| | | Low | High |
| Disease status | Localized | 63 | 56 |
| | Metastatic | 3 | 3 |

*P* = 0.888
Relative risk = 1.059
