## Supplementary figures and images for "Functional spatial transcriptomics uncover LMO7 as a fusion-regulated and clinically relevant driver of metastasis in Ewing sarcoma"

### Supplementary Figure 2

**Supplementary Figure 2.** Bursic *et al.*

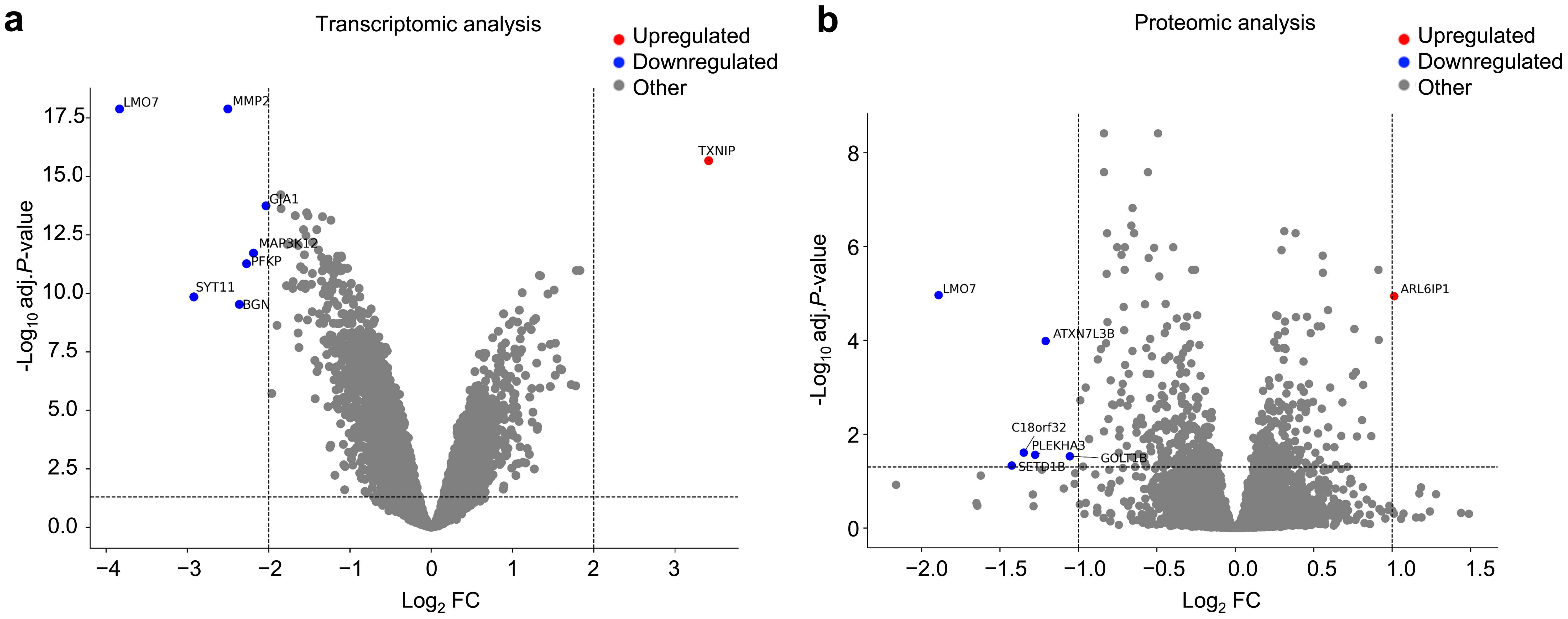

### Supplementary Figure 3

**Supplementary Figure 3.** Bursic *et al.*

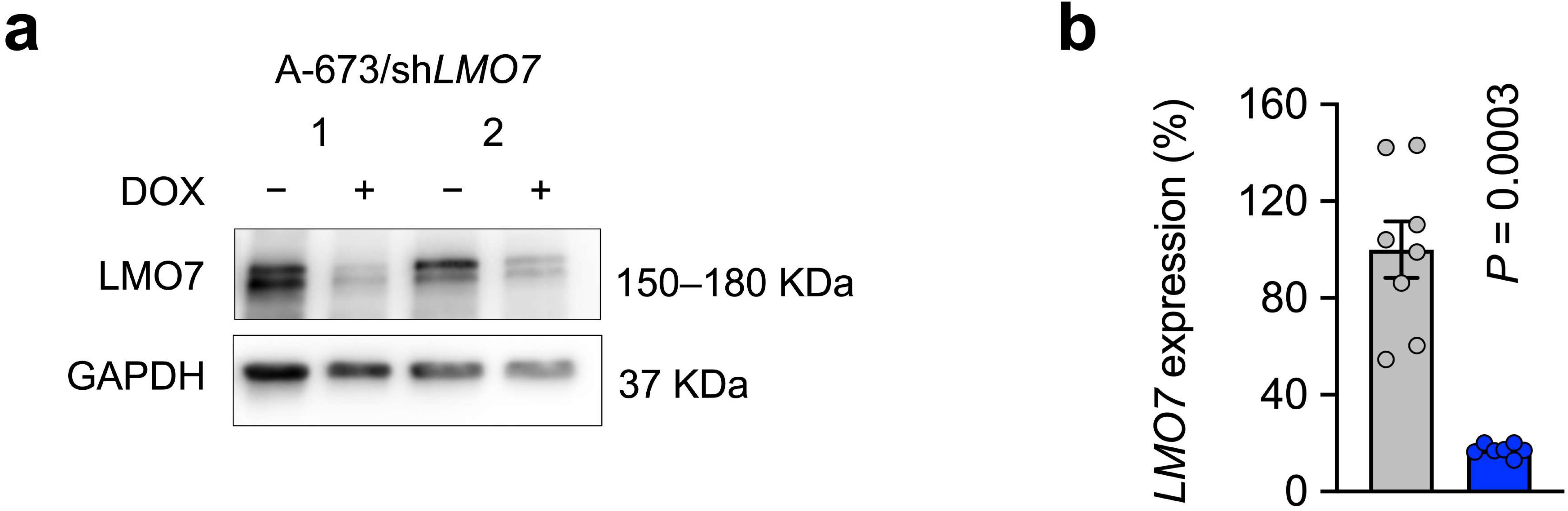
